# Impaired IL-10 Receptor Signaling Leads to Inflammation Induced Exhaustion in Hematopoietic Stem Cells

**DOI:** 10.1101/2025.09.30.679613

**Authors:** William Lucas Wadley, Helen Huang, Hew Yeng Lai, Jianhong Heidmann, Jane Chen, Eli Soyfer, Kalei Guillermo, Eshika Arora, Lauren Chen, Brianna Hoover, Angela Fleischman

## Abstract

Hematopoietic stem cells require tight regulation to rapidly initiate emergency hematopoiesis in response to pathogens, but chronic activation leads to proliferation induced exhaustion. Timely reentry into quiescence after inflammatory stimuli is essential for long term sustained HSC maintenance. We identify IL-10R signaling, an established negative feedback regulator in mature myeloid cells, as critical for returning HSCs to quiescence. IL-10R blockade prolongs HSC cycling and sustains activated transcriptional programs after acute inflammation. With chronic exposure, blockade increases cumulative divisions and accelerates aging hallmarks, including myeloid bias, loss of polarity, and functional defects, under conditions that do not otherwise exhaust HSCs when IL-10R signaling is intact. *Jak2^V617F^* mutant HSCs resist the aging acceleration induced by blockade. Consistent with this resistance, IL-10R blocking antibody promotes *Jak2^V617F^* clonal expansion and augments the myeloproliferative neoplasm phenotype. Together, these findings identify IL-10R signaling as a key coordinator of post inflammatory return to quiescence and suggest that modulating this axis could preserve HSCs and shape clonal hematopoiesis.

**Summary:** Wadley et al. show that IL-10 receptor signaling restrains inflammation-induced hematopoietic stem cell cycling and exhaustion; its blockade prolongs cycling, accelerates aging-related decline, and selectively favors *Jak2^V617F^* mutant HSCs, establishing IL-10 signaling as a critical regulator of inflammatory HSC exhaustion and malignant clonal evolution.

## Introduction

Hematopoietic stem cells (HSCs) reside at the apex of the hematopoietic hierarchy, generating all mature blood lineages through differentiation into progressively restricted progenitor populations (Pietras et al., 2014). To preserve long-term function, HSCs are maintained in quiescence, which protects this pool from the damaging effects of excessive proliferation (Orford and Scadden, 2008). Quiescence is tightly regulated by intrinsic programs, including FoxO3a, p53, and cyclin-dependent kinase inhibitors, and by extrinsic cues such as TGF-beta and Notch (Dickson et al., 1995; Soma et al., 1996). Under homeostatic conditions, HSCs remain largely dormant but can rapidly activate in response to inflammatory stimuli, such as infection, to support emergency hematopoiesis (Wilson et al., 2008). A timely return to quiescence after activation is essential to prevent stem cell exhaustion, which occurs if cycling persists.

Many hallmarks of stem cell exhaustion, including impaired regenerative capacity, expansion of phenotypic HSCs with reduced function, increased inflammatory signaling, and a shift toward myeloid-biased differentiation, mirror features of aged hematopoiesis (Dellorusso et al., 2024). Aging is associated with chronic, low-grade inflammation that stresses the HSC pool, driving both functional decline and selective pressure for clones that resist inflammation-induced exhaustion (López-Otín et al., 2013). This pressure contributes to clonal hematopoiesis of indeterminate potential (CHIP) (Jaiswal et al., 2014), in which HSCs harboring mutations such as *TET2* or *DNMT3A* expand over time. Other mutations, such as *PPM1D*, appear to be selected under specific stressors, for example chemotherapy exposure (Bolton et al., 2019; Hsu et al., 2018; Kahn et al., 2018). In contrast, *JAK2^V617F^*-driven clonal hematopoiesis, which can progress to overt myeloproliferative neoplasm (MPN), is relatively uncommon in the general aging population yet enriched in familial cases (Landgren et al., 2008; Ranjan et al., 2012). This pattern suggests shared environmental or inflammatory factors within MPN families that may preferentially favor *JAK2^V617F^*clone expansion.

Although much is known about maintaining HSC quiescence under steady-state conditions, the signals that govern the return to quiescence after inflammatory activation remain poorly defined. IL-10 is an anti-inflammatory cytokine that restrains excessive immune activation by suppressing responses downstream of Toll-like receptor (TLR) stimulation. In monocytes, IL-10 functions in a negative feedback loop to limit pro-inflammatory cytokine production (Saraiva and O’Garra, 2010). Mice lacking IL-10 exhibit defects in HSC function and recovery (Kang et al., 2007). Together, these findings suggest that IL-10 and its receptor may play a broader role in regulating responses to inflammation in HSC just as it does in monocytes.

Disruption of IL-10 receptor (IL-10R) signaling has been linked to severe inflammatory disorders and autoimmunity in humans and mice (Grundtner et al., 2009; Kennedy et al., 2000; Kuhn et al., 1993; Saraiva and O’Garra, 2010). However, more subtle defects in IL-10R signaling may permit persistent inflammation without overt autoimmunity. We previously showed that primary monocytes from patients with MPNs exhibit defective IL-10R signaling, resulting in prolonged TNF production following TLR stimulation (Lai et al., 2019). Notably, the same defect was observed in the unaffected identical twin of an MPN patient, suggesting a heritable or predisposing impairment in IL-10-mediated inflammatory resolution.

MPNs are clonal hematologic malignancies characterized by increased myelopoiesis, chronic inflammation, and association with aging (Campbell and Green, 2006; Fleischman, 2015; Soyfer and Fleischman, 2023). The *JAK2^V617F^*mutation, the most common driver in MPNs, promotes cytokine production and myeloid proliferation. However, in murine models, *Jak2^V617F^* does not confer a clear competitive advantage over wild-type HSCs (Mullally et al., 2010) and can impair stem cell function in knock-in models (Castiglione et al., 2021; Li et al., 2010). These observations suggest that additional factors are required to drive clonal expansion in vivo. The finding that both mutant and wild-type monocytes from MPN patients exhibit persistent inflammatory signaling supports an altered inflammatory environment rather than a cell-intrinsic property of *JAK2^V617F^*-mutant cells.

In this study, we examined the role of IL-10R signaling in regulating HSC responses to acute and chronic inflammation. We show that disruption of IL-10R function impairs the return to quiescence after inflammatory activation, accelerates inflammation-induced exhaustion of HSCs, and promotes expansion of *Jak2^V617F^*-mutant clones. These findings identify IL-10R signaling as a key regulator of stem cell fitness during inflammatory stress and suggest that intact IL-10R signaling protects the HSC pool from inflammation-induced exhaustion and clonal outgrowth.

## Results

### IL-10R blockade prolongs cell cycling of HSCs after an acute inflammatory stimulus

First, we tested whether blockade of IL-10R signaling alters HSC cell cycle dynamics after an acute inflammatory stimulus. In wild-type (WT) mice, we administered a single dose of the Toll-like receptor 4 (TLR4) ligand lipopolysaccharide (LPS), an inflammatory stimulant, with or without an IL-10R-blocking antibody (αIL-10R). Bone marrow was collected 24 and 72 hours after exposure (Figure 1A); bromodeoxyuridine (BrdU) was additionally injected 16 hours before harvest to label proliferating cells (Figure 1A). At 24 hours, we observed a similar increase in the percentage of proliferating hematopoietic stem and progenitor cells (HSPCs; Lin^−^/c-Kit^+^/Sca-1^+^, “LKS”) and long-term HSCs (LT-HSCs; CD150^+^/CD48^−^ LKS, “LKS-SLAM”) (Pietras et al., 2015) in either the LPS and LPS + αIL-10R groups (Figure 1B, 1C) (Figure S1A, S1B). By 72 hours, the proportion of BrdU^+^ LT-HSCs and HSPCs in the LPS-alone group had returned to baseline, whereas in the LPS + αIL-10R group the proportion of BrdU^+^ LT-HSCs and HSPCs remained at or above 24-hour levels (Figure 1B, 1C). These data indicate that IL-10R blockade doesn’t augment the initial proliferative response to inflammation but instead sustains the proliferative response of hematopoietic stem and progenitor compartments following an acute stimulus.

**Figure 1.**
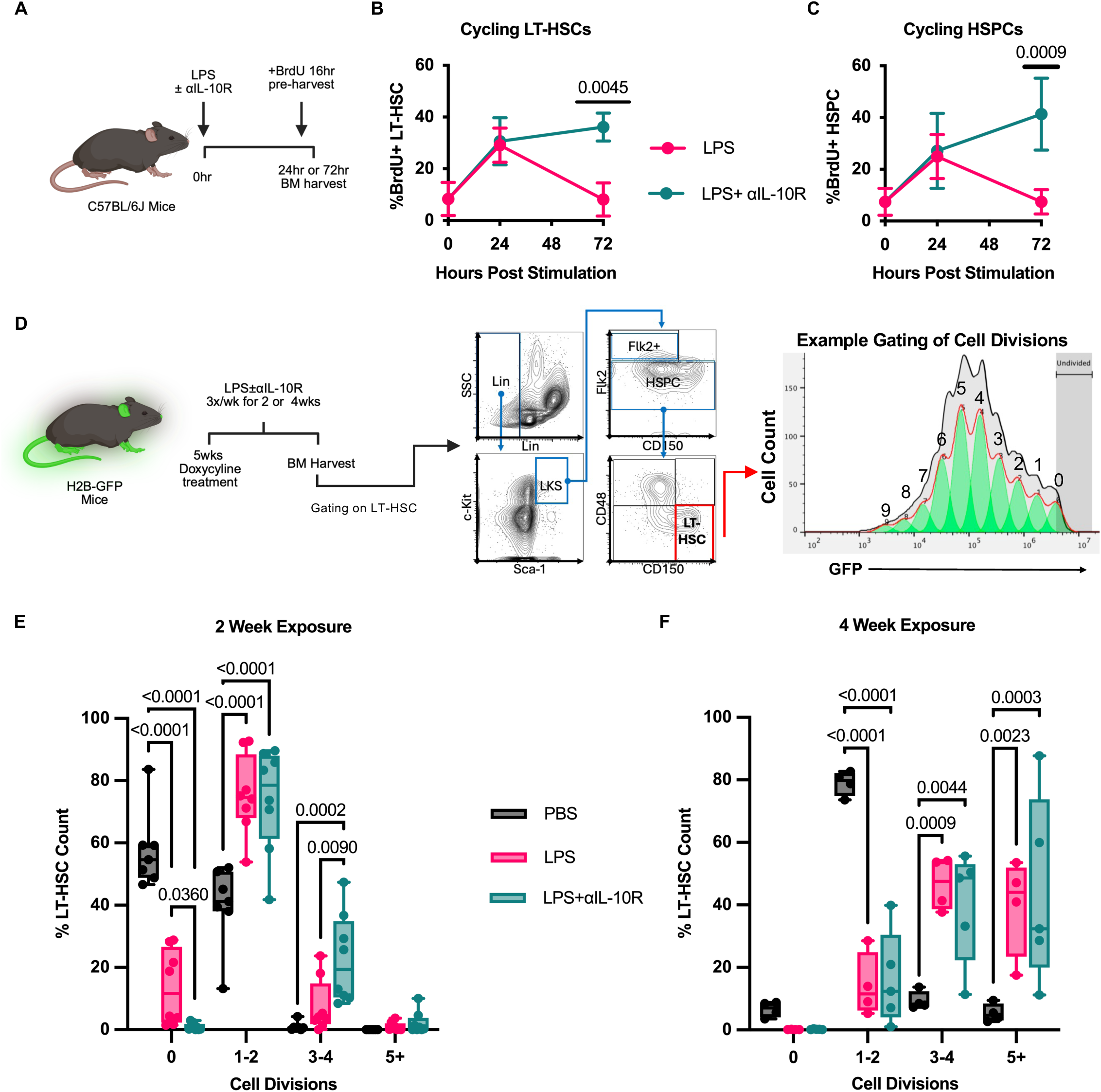
Impaired IL-10 signaling increases HSC proliferation. (A) Experimental model of acute inflammatory exposure and BrdU assay. (B) BrdU quantification of LT-HSCs from acute exposure with LPS and IL-10R blocking antibody, n = 3, SD error bars presented. (C) BrdU quantification of HSPCs from acute exposure with LPS and IL-10R blocking antibody, n = 3, SD error bars presented. (D) Experimental model of chronic inflammatory exposure using H2B-GFP^+^ mice (*left*), example gating scheme (*middle*) and example of H2B-GFP proliferative history (*right*). (F) Proliferative history of LT-HSCs in H2B-GFP mice exposed for 2 weeks to LPS (n = 8) with or without IL-10R blocking antibody (n = 8), or PBS as control (n = 7). All dots in box and whisker plot represent each data point. (F) Proliferative history of LT-HSCs in H2B-GFP mice exposed for 4 weeks to LPS (n = 4) with or without IL-10R blocking antibody (n = 4), or PBS as control (n = 5). All dots in box and whisker plot represent each data point. Statistical significance was determined by unpaired *t* test with Holm-*Śídák* correction (B and C) and ordinary two-way ANOVA with Tukey’s correction (E and F). If not specifically indicated, P-value < 0.05, 0.005, 0.0005, and <0.0001 are represented as *, **, ***, and **** respectively.

### IL-10R blockade increases proliferative history of HSCs during chronic inflammatory stress

We next examined the role of IL-10R signaling in regulating HSC proliferation under conditions of chronic inflammatory stress. To quantify proliferative history, we used a doxycycline-inducible H2B-GFP reporter mouse model in which hematopoietic cells, including HSCs, incorporate GFP during doxycycline exposure. Upon doxycycline withdrawal, dilution of GFP provides a readout of cumulative cell divisions (Foudi et al., 2009). We assessed long-term HSCs (LT-HSCs) after two or four weeks of chronic LPS exposure with or without IL-10R blockade (αIL-10R) (Figure 1D, Figure S1C). In PBS treated controls, most LT-HSCs underwent zero to two divisions, with only a small fraction dividing more extensively, consistent with rare HSC cycling under steady-state conditions (Figure 1E, 1F). In contrast, LPS-treated mice showed significantly increased proliferation, with LT-HSCs averaging one to two divisions after two weeks and approximately 40% of cells undergoing five or more divisions after four weeks (Figure 1E, 1F). At the four-week time point, both LPS and LPS + αIL-10R groups demonstrated marked proliferation, and no statistical differences were detected (Figure 1F). However, at two weeks, when the proliferative response to LPS alone remained moderate, LT-HSCs from the LPS + αIL-10R group exhibited a significantly greater number of divisions compared with LPS alone (Figure 1E). Collectively, these findings suggest that IL-10R blockade accelerates HSC cycling and predisposes to exhaustion when brief inflammatory exposures are encountered that would not otherwise provoke excessive proliferation.

### Pro-inflammatory pathways persist in the presence of IL-10R blockade

Because IL-10R blockade extended HSC cycling after an acute inflammatory stimulus, we asked whether transcriptomic programs also persist, complementing the phenotype. We again exposed mice to a single dose of LPS with or without αIL-10R at 24 and 72 hours, as described previous, and performed bulk RNA sequencing on flow cytometry sorted HSPCs (Figure 2A) (Figure S1A). Gene sets were defined and clustered using the MSigDB mouse collections, and RNA-seq data were processed with DESeq2 to identify differentially expressed genes (DEGs; p < 0.05) (Subramanian et al., 2005), we then generated heatmaps of significant gene sets and their respective DEGs. Single sample gene set enrichment analysis was performed on the RNA-seq data to observe pathway activation across samples by way of gene set variation analysis (GSVA) and gene set enrichment analysis (GSEA) (Subramanian et al., 2005). At 24 hours post exposure, both LPS and LPS + αIL-10R groups showed a marked change in gene expression signatures relative to PBS, indicating a robust inflammatory and proliferative response (Figure 2B, 2D) (Figure S2). By 72 hours, MSigDB gene-set gene expression in HSPCs after the LPS exposure returned toward baseline and resembled PBS controls, whereas in the LPS + αIL-10R group the initial inflammatory signatures are continued (Figure 2B, 2D). Gene sets linked to DNA replication and DNA repair were strongly induced at 24 hours and remained elevated only with αIL-10R, consistent with the previous sustained cycling data (Figure 1C) (Figure 2B, 2D). Similarly, pathways indexing inflammatory and metabolic activity, including E2F targets, oxidative phosphorylation, mTOR signaling, and the interferon-α response, were upregulated after LPS and, unlike LPS alone, remained elevated with αIL-10R (Figure 2B, 2D). In contrast, genes associated with HSC homeostasis were downregulated after LPS, returned to baseline by 72 hours in LPS alone, and remained suppressed with αIL-10R (Figure 2B). Principal component analysis (PCA) of the transcriptomic profiles revealed that samples from the PBS day-0 control group clustered closely with those collected 72 hours after LPS exposure along the main axis of variation (PC1), indicating transcriptional recovery towards baseline following the initial inflammatory insult. (Figure 2C) (Figure S2). In contrast, all other groups remain transcriptionally distinct and cluster around themselves, reflecting the immediate inflammatory response and the prolonged response when paired with impaired IL-10R signaling (Figure 2C). Together, these data show that impaired IL-10R signaling sustains acute inflammation–induced transcriptional programs in HSPCs, mirroring the prolonged cycling we observe at 72 hours under the same conditions (Figure 1C).

**Figure 2.**
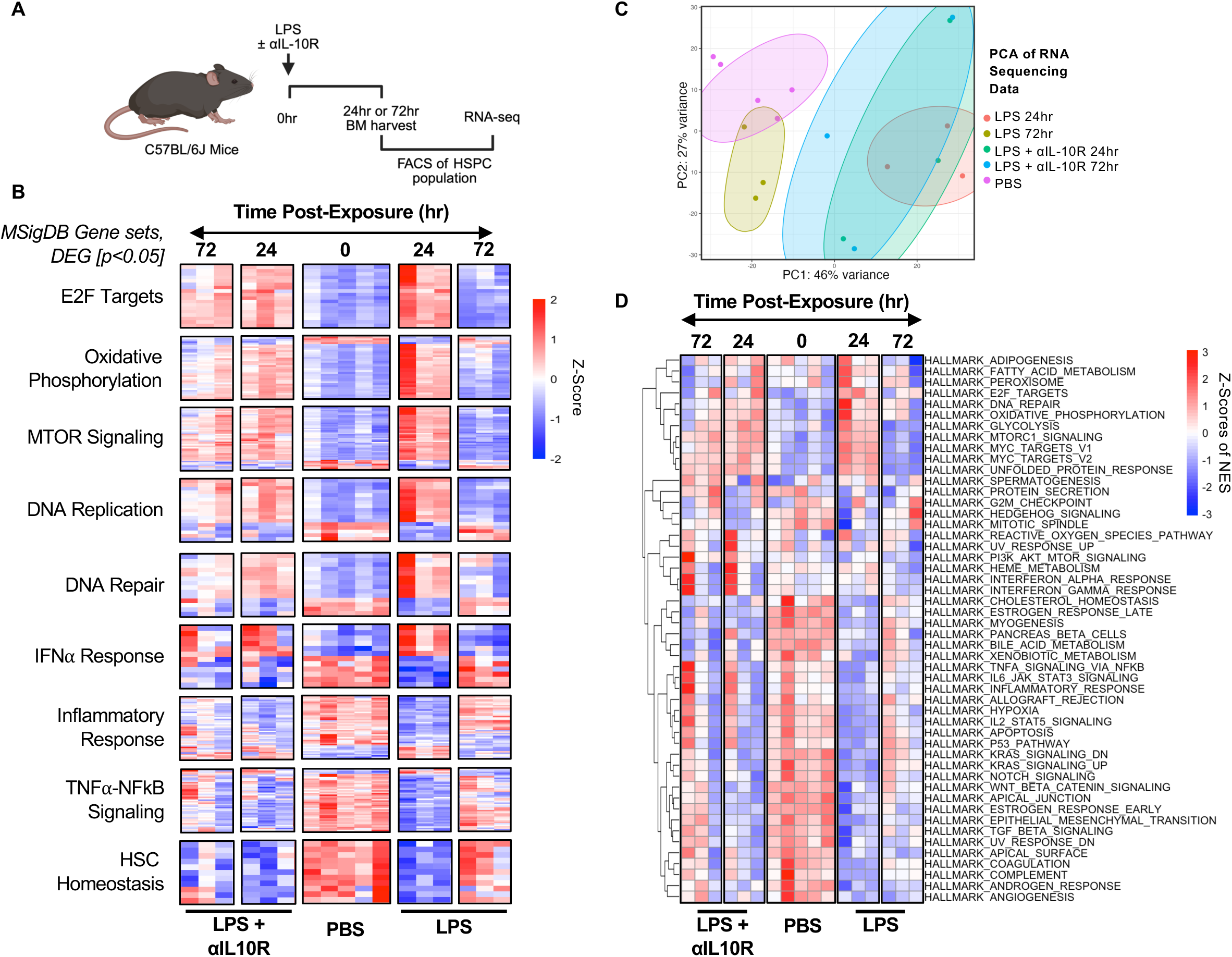
Inflammatory gene expression signatures are prolonged with impaired IL-10R signaling. (A) Experimental model of acute inflammatory exposure and bulk RNA-seq with FAC sorted HSPCs. (B) Heat map clusters of various MSigDB gene set associated differentially expressed genes with a p-value of less than 0.05, highlighting gene expression signatures in between exposure conditions. (C) Principal component analysis (PCA) of the bulk-cell RNA-seq data. (D) Single sample gene set enrichment analysis (ssGSEA) for pathway enrichment across samples and groups. Statistical significance was determined by Wald test with Benjamini–Hochberg FDR correction (B) as standard using DESeq2.

### Impaired IL-10R signaling promotes a myeloid progenitor bias under inflammatory stress

Because myeloid bias is a hallmark of hematopoietic aging, we tested whether chronic IL-10R blockade in the context of TLR4 stimulation (LPS) alters the balance of myeloid- and lymphoid-restricted progenitors (Figure 3A,B). The overall frequency of Lin⁻/c-Kit⁺/Sca-1⁻ (LK) cells, representing myeloid-restricted progenitors (MRPs), was unchanged by LPS or LPS + αIL-10R (Figure 3C, Figure S1D) (Challen et al., 2009; Cozzio et al., 2003). However, analysis of defined subsets revealed significant shifts. After four weeks of LPS or LPS + αIL-10R exposure, common myeloid progenitors (CMPs; CD34^high^/CD16/32^low^) were reduced, with a reciprocal increase in granulocyte–macrophage progenitors (GMPs; CD34^+^/CD16/32^+^), while megakaryocyte erythroid progenitors (MEPs; CD34^−^/CD16/32^low^) remained stable (Figure 3E, Figure S1D). Notably, GMP expansion was already evident after two weeks in the LPS + αIL-10R group, but not in LPS alone, consistent with the model that impaired IL-10R signaling accelerates hematopoietic aging (Figure 3E).

**Figure 3.**
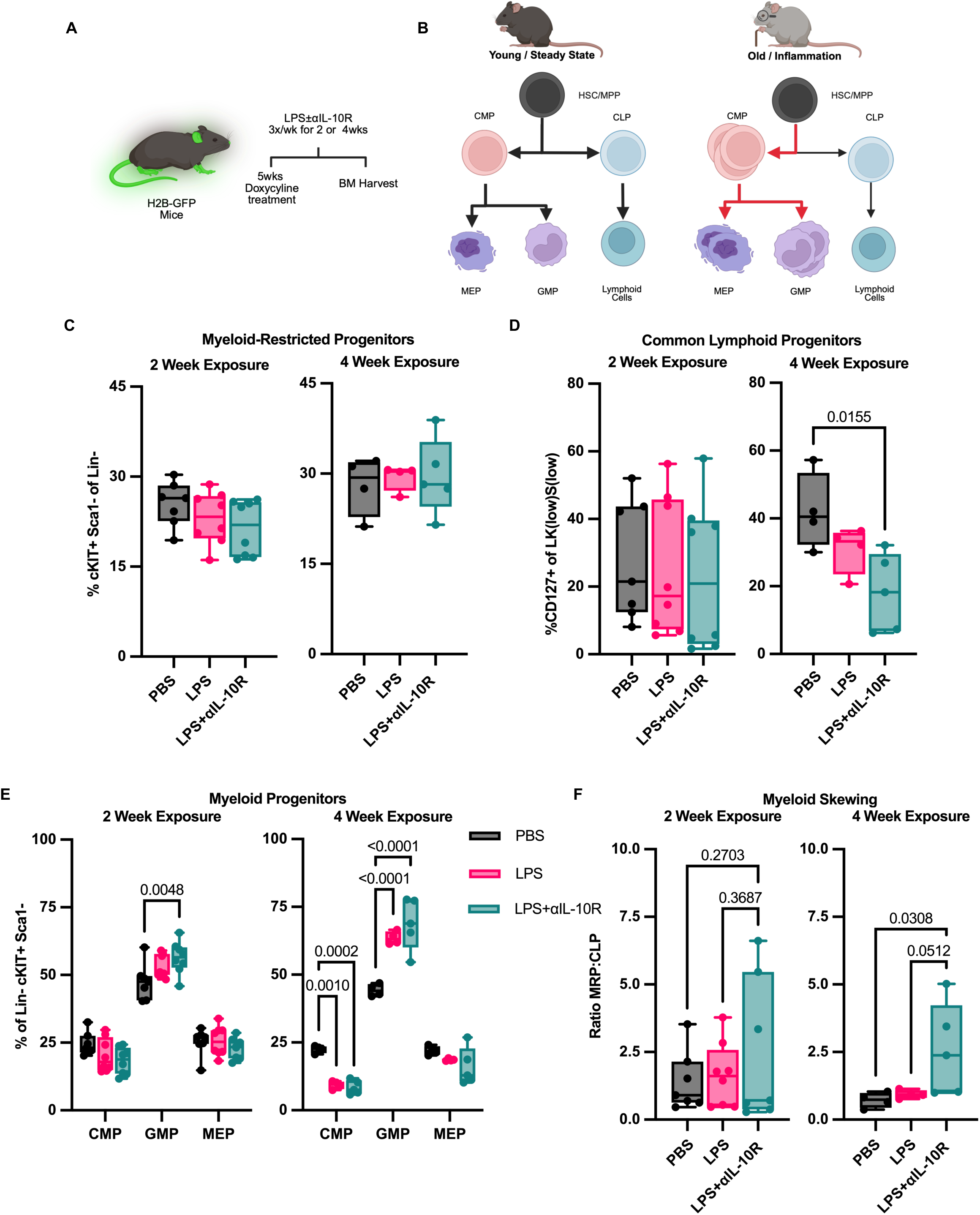
Aged phenotype of myeloid skewing is hastened with impaired IL-10R signaling. (A) Experimental model of chronic inflammatory exposure using H2B-GFP mice model. (B) Graphical abstract of myeloid skewing from aging or inflammation. (C) Frequency of myeloid-restricted progenitors (MRP) in lineage negative population after 2 weeks (*left*) exposed to LPS (n = 8) with or without IL-10R blocking antibody (n = 8), or PBS as control (n = 7); or 4 weeks (*right*) exposed to LPS (n = 4) with or without IL-10R blocking antibody (n = 4), or PBS as control (n = 5). (D) Frequency of common lymphoid progenitors (CLP) after 2 weeks (*left*) or 4 weeks (*right*). (E) Sub-population composition within MRP group, consisting of common myeloid progenitor (CMP), granulocyte monocyte progenitor (GMP), and megakaryocyte erythroid progenitor (MEP). After 2 weeks (*left*) or 4 weeks (*right*). (F) Graphed myeloid skewing representation through the ratio of MRP to CLP populations after 2 weeks (*left*) or 4 weeks (*right*). All dots in box and whisker plot represent each data point. Statistical significance was determined by ordinary one-way ANOVA (C, D, and F) with Tukey’s correction, and ordinary two-way ANOVA (E) with Tukey’s correction. If not specifically indicated, P-value < 0.05, 0.005, 0.0005, and <0.0001 are represented as *, **, ***, and **** respectively.

Because increased myeloid output without loss of lymphoid potential does not constitute exhaustion, we next assessed common lymphoid progenitors (CLPs; Lin⁻/c-Kit^low^/Sca-1^low^/CD127⁺). CLPs were significantly reduced only in the LPS + αIL-10R group after four weeks, but not at two weeks and not with LPS alone, indicating that impaired IL-10R signaling accelerates lymphoid decline (Figure 3D, Figure S1D). This lineage imbalance was further reflected in the ratio of myeloid-restricted to lymphoid progenitors, which shifted toward myeloid dominance only after four weeks of LPS + αIL-10R (Figure 3F). Thus, although total myeloid progenitor frequency remained stable, the selective increase in GMPs combined with reduction in CLPs establishes a functional myeloid bias that emerges under prolonged inflammatory stress when IL-10R signaling is impaired.

### Chronic IL-10R Blockade Reduces Reconstitution Capacity in the Setting of Chronic Inflammation

Loss of polarity in HSPCs is a feature of aging and is associated with impaired migration and functional decline (Amoah et al., 2022; Florian et al., 2012). We therefore initially assessed the impact of chronic LPS with or without αIL-10R on HSPC polarity of alpha-tubulin using image flow cytometry (IFC) (Figure 4A) (Figure S1A). Image flow cytometry showed a reduction in polar HSPCs from 50% in controls to 35% after LPS. This effect was amplified by IL-10R blockade, with ∼25% of cells exhibiting polarity, which corresponds to a ∼50% decrease relative to controls (Figure 4B) (Jespersen et al., 2023).

**Figure 4.**
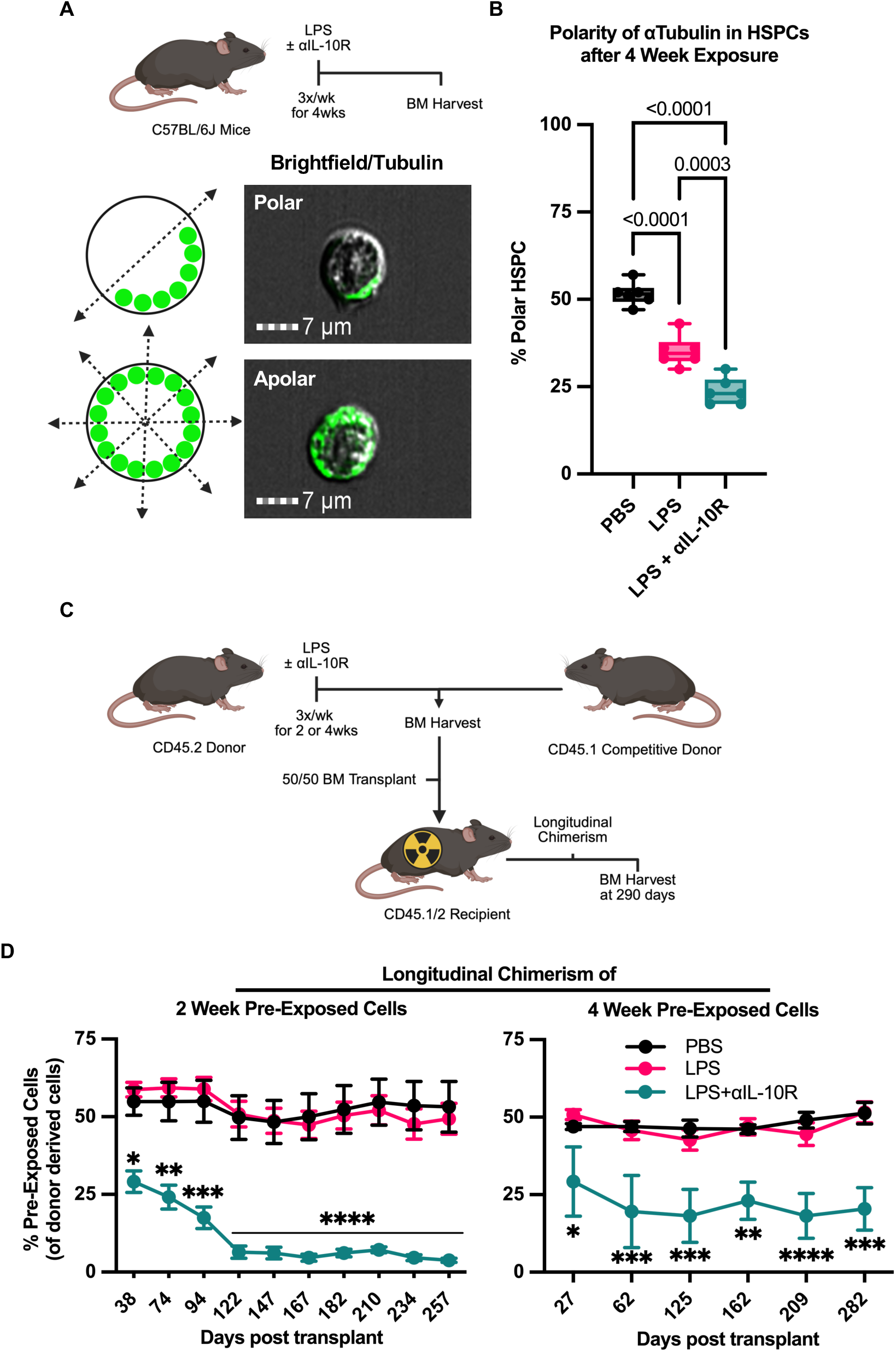
Impaired IL-10 signaling reduces reconstitution capacity during competitive transplantation after inflammatory stress. (A) Experimental model of chronic inflammatory exposure for polarity assay. (B) Alpha-tubulin polarity of HSPCs after chronic exposure to inflammatory stress. PBS control (n = 6), LPS (n = 6), or LPS with anti-IL-10R blocking antibody (n = 6). (C) Experimental model for chronic inflammatory pre-exposure followed by competitive transplantation in irradiated host mice. (D) Longitudinal chimerism post-transplant of donor mice exposed to inflammatory stress for 2 weeks (*left*) with PBS control (n = 10), LPS (n = 10), and LPS with IL-10R blocking antibody (n = 5). Or for 4 weeks (*right*) with PBS control (n = 6), LPS (n = 8), and LPS with IL-10R blocking antibody (n = 6). All statistical comparisons between PBS and LPS groups in either time group were non-significant. SEM error bars presented. All dots in box and whisker plot represent each data point. Statistical significance was determined by ordinary one-way ANOVA (B) with Tukey’s correction, and repeated measures (RM) two-way ANOVA (D) with *Śídák* correction. If not specifically indicated, P-value < 0.05, 0.005, 0.0005, and <0.0001 are represented as *, **, ***, and **** respectively.

To assess function, we performed competitive transplantation. CD45.2 donor mice received PBS, LPS alone, or LPS + αIL-10R three times weekly for two or four weeks. Bone marrow (BM) from treated donors was mixed 1:1 with untreated, age- and sex-matched CD45.1 BM and transplanted into lethally irradiated CD45.1/2 recipients (Figure 4C). Donor-derived hematopoietic reconstitution was monitored longitudinally in peripheral blood (Figure S3A). Recipients of BM from the LPS + αIL-10R group exhibited significantly impaired repopulation compared with PBS or LPS alone, in either time group, with progressive loss of donor chimerism over time (Figure 4D). To directly assess reconstitution at the single HSC level after two weeks of LPS or LPS + αIL-10R, we treated mice, sorted LT HSCs, and established competitive transplants mixing 250 PBS treated LT HSCs with 250 LT HSCs from LPS + αIL-10R donors. Mirroring the whole bone marrow results, LT HSCs from LPS + αIL-10R donors performed inferiorly to PBS treated LT HSCs (Figure S3C). Additional mice were treated with αIL-10R alone for four weeks prior to transplantation to observe the impact of IL-10R blockade in the absence of additional inflammatory stimuli on competitive transplant fitness. Competitive repopulation did not differ from controls (Figure S3B), indicating that IL-10R blockade in the absence of inflammatory stimulation does not appreciable impair reconstitution capacity. Together these findings demonstrate that chronic IL-10R blockade during inflammatory stress compromises HSC function, reflected by reduced polarity and consequential diminished reconstitution capacity.

### IL-10R Blockade Does Not Induce Prolonged Cycling in Jak2^V617F^ HSPCs

We previously found that monocytes from patients with myeloproliferative neoplasms (MPNs) exhibit impaired IL-10R signaling (Lai et al., 2019). This defect was also present in an unaffected identical twin of an MPN patient but absent in *Jak2^V617F^* mice, suggesting that impaired IL-10R signaling may reflect a germline susceptibility trait rather than a consequence of the somatic *JAK2^V617F^* mutation. We therefore hypothesized that *JAK2^V617F^* renders HSCs less responsive to the proliferative effects of IL-10R blockade, conferring a selective advantage to mutant HSCs under defective IL-10R signaling.

To test this hypothesis, we compared LT-HSC proliferation using BrdU incorporation. We administered a single acute dose of LPS with or without αIL-10R to *Jak2^V617F^* mice and evaluated responses alongside wild-type (WT) controls (Figure 5A) (Figure S1B). In WT mice, LT-HSC proliferation increased at day 1 after LPS and declined by day 3. Addition of αIL-10R prolonged this proliferative response, consistent with earlier results (Figure 5B) (Figure 1B). In contrast, *Jak2^V617F^* LT-HSCs displayed elevated basal cycling relative to WT but showed minimal additional proliferation after LPS (Figure 5C). Notably, αIL-10R did not augment this response and modestly reduced BrdU incorporation by day 3 (Figure 5C). Similar patterns were observed in the broader HSPC compartment (Figure S4A, S4B).

**Figure 5.**
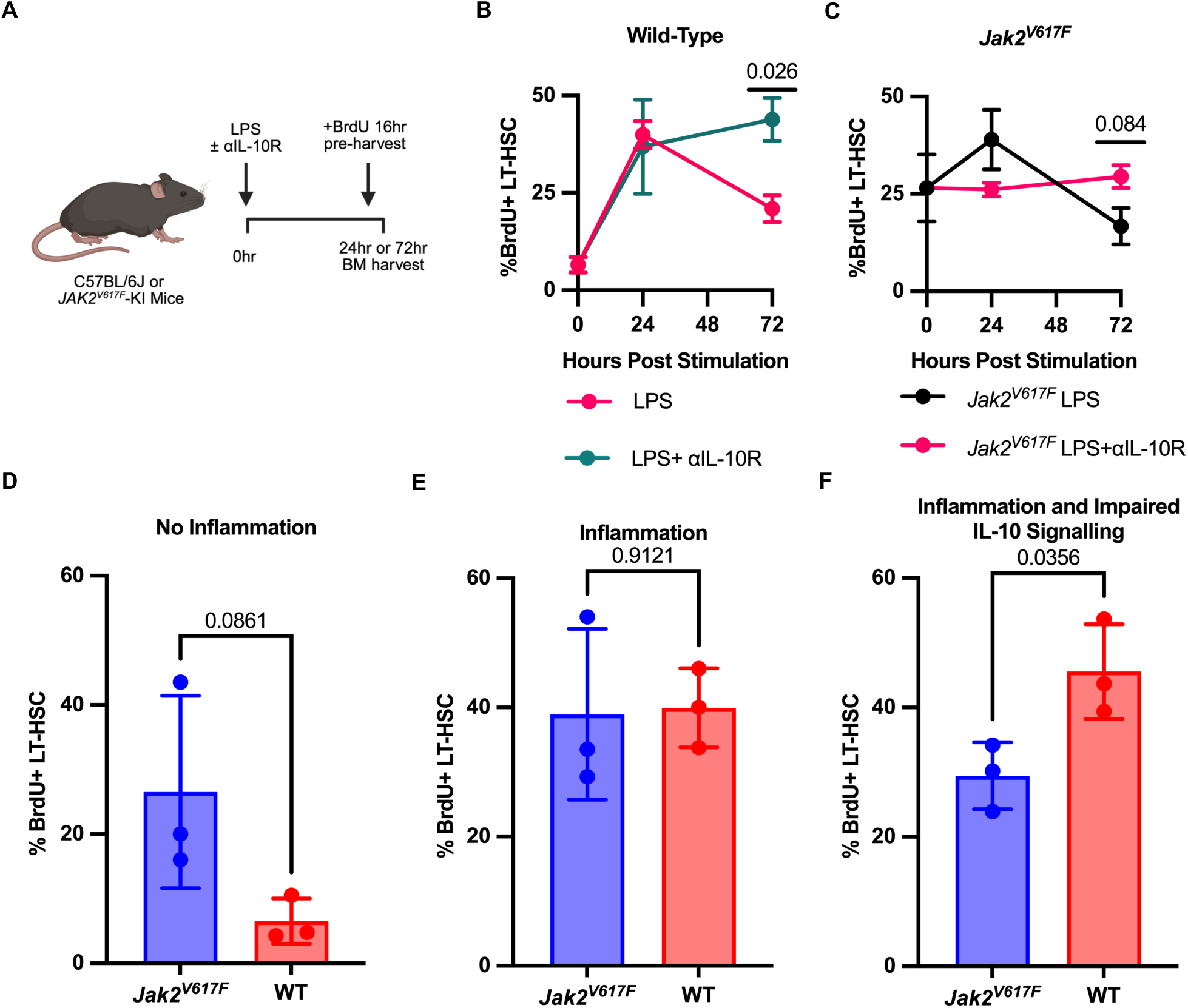
*Jak2^V617F^* LT-HSC cycling is not affected by impaired IL-10R signaling in presence of acute inflammatory stress. (A) Experimental model of acute inflammatory exposure and BrdU assay with WT or *Jak2^V617F^* mutant mice. (B) BrdU quantification of WT LT-HSCs from acute exposure with LPS and IL-10R blocking antibody (n = 3 for all conditions and groups), SD error bars presented. (C) BrdU quantification of *Jak2^V617F^* LT-HSCs (*right*) from acute exposure with LPS and IL-10R blocking antibody (n = 3 for all conditions and groups), SD error bars presented. (D-F) Graphical summary of LT-HSC cycling under each condition between WT and *Jak2^V617F^* mutants. All dots in bar plots represent a biological replicate data point. Statistical significance was determined by unpaired *t* test with Holm-*Śídák* correction (B and C), and two-tailed unpaired *t* test (D, E, and F). If not specifically indicated, P-value < 0.05, 0.005, 0.0005, and <0.0001 are represented as *, **, ***, and **** respectively.

These findings indicate that *Jak2^V617F^* LT-HSCs have elevated baseline cycling and reduced sensitivity to IL-10 dependent regulation of inflammation-induced proliferation compared with WT LT-HSCs (Figure 5D). Following inflammatory stimulation, both WT and *Jak2^V617F^* LT-HSCs enter the cell cycle to a similar degree (Figure 5E). However, in the context of IL-10R blockade, WT LT-HSCs show sustained cycling, whereas *Jak2^V617F^*LT-HSCs do not (Figure 5F). This blunted response may protect mutant HSCs from inflammation-driven proliferative exhaustion and preserve long-term function, thereby conferring a relative fitness advantage over time.

### Chronic IL-10R Blockade Reveals a Selective Advantage of Jak2^V617F^ HSPCs

To test the functional impact of IL-10R signaling on the long-term competitive fitness of *Jak2^V617F^*-mutant versus WT HSCs, we performed competitive transplantation using equal ratios of HSC-equivalent whole bone marrow from *Jak2^V617F^* or WT donors (CD45.2) co-transplanted with CD45.1 WT competitor cells into lethally irradiated CD45.1/2 recipients (Figure 6A). Beginning 60 days post-transplantation, mice were either treated with an IL-10R–blocking antibody or PBS control. In the PBS treated *Jak2^V617F^*group, the contribution of *Jak2^V617F^*-derived peripheral blood cells progressively declined over time (Figure 6B). In contrast, IL-10R blockade significantly increased *Jak2^V617F^* peripheral blood chimerism by day 60 after the start of the treatment, compared to PBS treated mice (Figure 6B). As a control, IL-10R blockade had no effect on WT:WT competitive transplants (Figure S4C).

**Figure 6.**
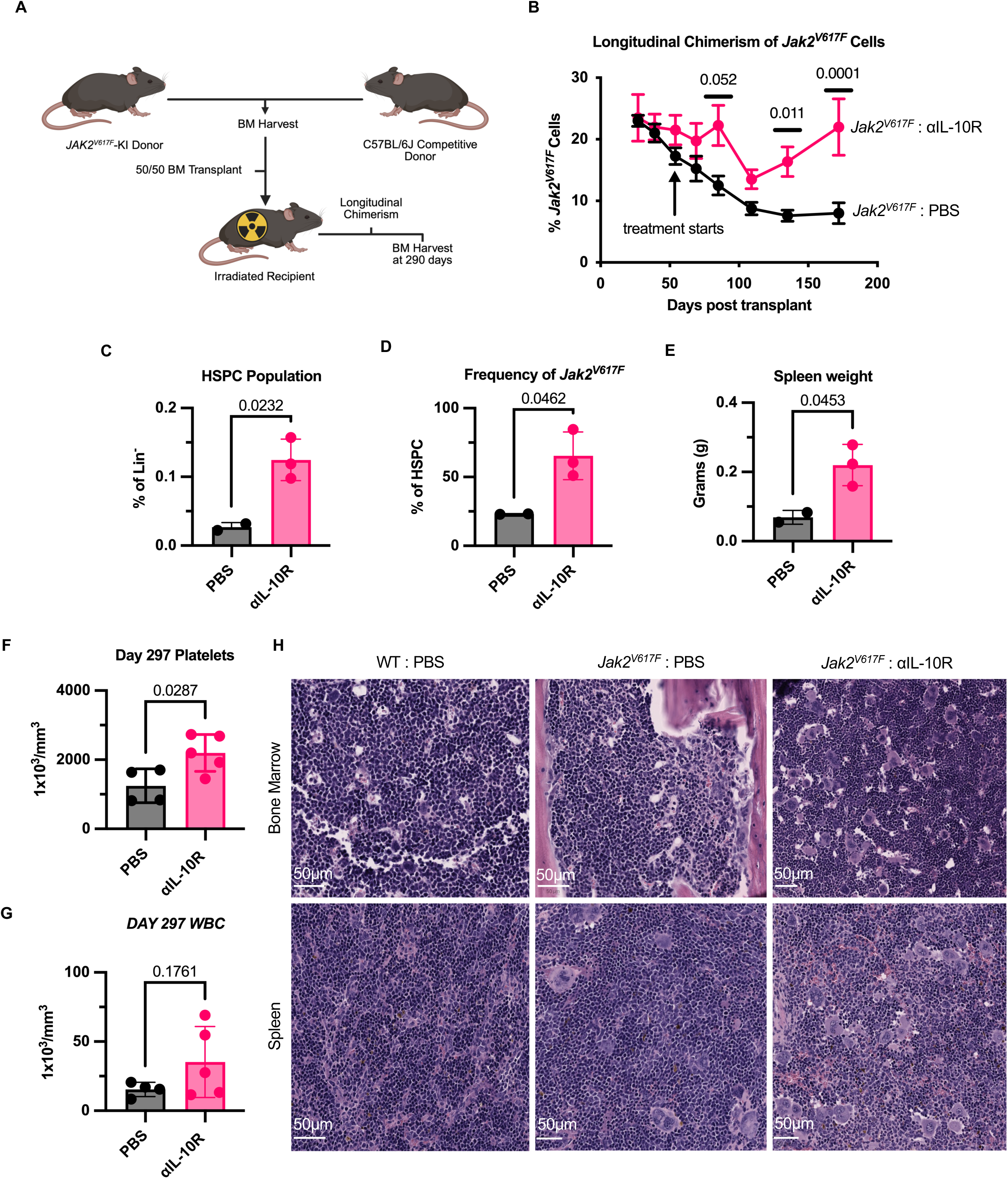
Impaired IL-10R signaling allows MPN disease pathology from *Jak2^V617F^*. (A) Experimental model of competitive transplantation with WT of *Jak2^V617F^*mutant mice. (B) *Jak2^V617F^* chimerism and starting a blocking IL-10R antibody treatment (n = 5) 60-days post-transplantation, or no treatment control (n = 4). SEM error bars presented. (C) HSPC quantification at end of transplant comparing the aIL-10R treatment (n = 3) and control (n = 2). (D) Quantification of *Jak2^V617F^*+ in the HSPC population after treatment. (E) Spleen weight of transplant mice after treatment. (F) Quantification of platelets at end of transplant with aIL-10R treatment (n = 5), and control (n = 4). (G) Quantification of white blood cells at end of transplant with aIL-10R treatment (n = 5), and control (n = 4). (H) Histology of bone marrow (*top*) and spleen (*bottom*) indicating megakaryocyte infiltration in *Jak2^V617F^* transplant mice treated with aIL-10R. All bar plots present with SD error bars. All dots in bar plots represent a biological replicate data point. Statistical significance was determined by repeated measures (RM) two-way ANOVA with *Śídák* correction (B), and two-tailed unpaired *t* test (C - G). If not specifically indicated, P-value < 0.05, 0.005, 0.0005, and <0.0001 are represented as *, **, ***, and **** respectively.

To more directly test whether exposure to LPS plus αIL-10R preferentially impairs wild type relative to *Jak2^V617F^* HSC fitness, we treated *Jak2^V617F^* knock in and wild type donors with PBS or with LPS + αIL-10R for two weeks. We then sorted LT HSCs and competitively transplanted 250 wild type and 250 *Jak2^V617F^* LT HSCs from donors that had received the same treatment. When both donor populations were PBS treated, *Jak2^V617F^* LT HSCs were less competitive than wild type (Figure S3D). In contrast, when both donor populations were exposed to LPS + αIL-10R, *Jak2^V617F^* LT HSCs maintained the ability to compete with wild-type LT HSC (Figure S3E). These results support a model in which IL-10R blockade selectively compromises wild type HSCs, thereby increasing the relative fitness of *Jak2^V617F^*HSCs.

At sacrifice, bone marrow showed a higher frequency of HSPCs with a disproportionately greater contribution from *Jak2^V617F^* donor-derived cells in αIL-10R treated mice compared with PBS controls (Figure 6C, 6D) (Figure S1A) (Figure S3A). Spleen weights were increased in the αIL-10R cohort, consistent with a *Jak2^V617F^*-driven myeloproliferative phenotype (Fig. 6E). These transplantation results align with the proliferation data. *Jak2^V617F^* HSCs show elevated basal cycling and are therefore more prone to proliferative exhaustion, which makes them less competitive than wild type HSCs. When IL-10R signaling is blocked in an inflammatory milieu driven by the *Jak2^V617F^* clone, wild type HSCs undergo sustained cycling and exhaustion, whereas *Jak2^V617F^* HSCs are relatively spared, resulting in a selective advantage for the mutant clone over time.

### IL-10R blockade exacerbates MPN phenotype

We repeated the competitive transplantation of *Jak2^V617F^* and wild type bone marrow to assess how IL-10R blockade influences classical features of MPN in the *Jak2^V617F^* knock-in model, including thrombocytosis, splenomegaly, and megakaryocytic expansion in bone marrow. Mice received αIL-10R from day 49 after transplant through day 291 (Figure S4D). Mice received αIL-10R from day 49 after transplant through day 291 (Figure S4D). Consistent with prior findings, *Jak2^V617F^* chimerism expanded only during αIL-10R treatment (Figure S4D). αIL-10R–treated mice showed significantly higher platelet counts and a trend toward increased leukocyte counts (Figure 6F, 6G), indicating that IL-10R blockade not only provides a selective advantage to the mutant clone but also augments overt MPN features. At sacrifice, hematoxylin and eosin staining demonstrated increased megakaryocytes in bone marrow and prominent megakaryocytic infiltration of the spleen in αIL-10R–treated *Jak2^V617F^* mice (Fig. 6H). Collectively, these data show that chronic impairment of IL-10R signaling promotes *Jak2^V617F^* clonal expansion and exacerbates disease related phenotypes.

## Discussion

We identify IL-10R signaling as critical regulator that restores hematopoietic stem cell quiescence after inflammatory activation. Following acute LPS exposure, IL-10R blockade did not amplify the initial proliferative surge of LT-HSCs and HSPCs but prevented return to quiescence, as shown by persistent BrdU incorporation. Principal component analysis placed LPS exposed HSPCs near baseline by 72 hours, whereas LPS + αIL-10R HSPCs remained transcriptionally displaced. When this resolving signal is disabled, HSCs persist in a metabolically active cycling state that sustains inflammation-associated transcriptional programs and accelerates functional decline. These effects coincide with lineage skewing within progenitor compartments, loss of polarity, and diminished competitive repopulation, indicating that IL-10R signaling promotes reentry into quiescence and preserves HSC fitness after inflammatory activation. Mechanistically, IL-10 limits cytokine production in monocytes through a well-characterized negative feedback loop after TLR stimulation (de Waal Malefyt et al., 1991). Prior studies have implicated IL-10 in HSC fitness (Kang et al., 2007; McCabe et al., 2018), our data align with this framework and support a model in which IL-10R signaling terminates inflammation-induced cycling in HSCs.

Inherited or acquired defects in IL-10R signaling generate an inflammatory milieu that compromises wild-type HSC fitness while favoring expansion of inflammation-resistant clones such as *JAK2^V617F^*. Therapeutically, strategies that enhance inflammatory resolution by augmenting IL-10R signaling, modulating downstream effectors that normalize cell-cycle exit and metabolic activity, or targeting niche sources or targets of IL-10 may protect the HSC pool during chronic inflammatory states and potentially restrain clonal selection in high-risk settings. Conversely, medications or comorbid inflammatory conditions that blunt IL-10R signaling may accelerate HSC exhaustion and promote expansion of clones with mutations unresponsive to IL-10R blockade or resistant to cycling-induced attrition.

Several caveats frame interpretation. First, LPS engages TLR4 and models a bacterial stimulus, while informative it captures only part of the inflammatory spectrum relevant to aging and MPN. Different stimulus may elicit distinct IL-10R dependencies. Second, we used systemic antibody blockade rather than HSC-restricted genetic perturbation, thus HSC extrinsic effects, for example on monocytes and niche macrophages may likely contribute. Dissecting HSC-intrinsic vs. niche-mediated IL-10R actions will require conditional deletions and controlled reconstitution experiments. Third, single cell RNA-sequencing would refine the cellular populations and targets of HSC activity resolution. Finally, while we demonstrate a selective advantage of *JAK2^V617F^*clones, whether HSC bearing other more common CHIP mutations similarly benefit from IL-10R impairment remains to be tested directly. Prospective human studies to define the prevalence, penetrance, and molecular basis of IL-10R pathway defects in MPN families, and interventional studies testing IL-10 pathway augmentation, are warranted.

We propose that IL-10R signaling is a key arbiter of HSC activation during inflammation and of its resolution. Intact IL-10R enables timely quiescence re-entry, preserves stem cell fitness, and maintains balanced lineage output. When this axis is compromised, HSCs fail to disengage from cycling, accumulate replicative and metabolic stress, and yield competitive ground to inflammation-resistant clones such as *JAK2^V617F^*thereby granting clonal selection and progression of MPN disease phenotypes. These insights elevate inflammatory resolution as a central determinant of stem cell longevity and clonal evolution in the hematopoietic system.

## Methods

### Mice and genotyping

Wild-type C57BL/6J mice purchased from the Jackson laboratory (#000664). *Jak2^V617F^* ^fl/+^ mice (Mullally et al., 2010) were a gift from Ann Mullally. *Jak2^V617F^* ^fl/+^ mice were mated with Vav-Cre mice purchased from Jackson Laboratory (#008610) to create *Jak2^V617F^* knock-in mice. Genotyping of mice was performed with genomic DNA extracted from peripheral blood leukocytes using 30–50 µL of blood collected into 100 mM EDTA to prevent clotting, followed by erythrocyte lysis in ammonium chloride potassium (ACK) buffer. Pellets were then resuspended in lysis buffer (0.1 M Tris pH 8, 0.2 M NaCl, 5 mM EDTA, 0.4% SDS) containing proteinase K and incubated at 56°C for 2hr or overnight. DNA was precipitated by adding 100% isopropanol and centrifuging at 12,000 g for 3 min, followed by washing in 70% ethanol and air-drying for 10 min. The DNA pellet was resuspended in TE buffer and incubated at 56°C for 1 h before storage at −20°C or until use. PCR amplification was performed using DreamTaq Green DNA polymerase (EP0712; ThermoFisher Scientific) with primer sets specific for the Jak2V617F knock-in allele (J2KI-1: 5′-CGTGCATAGTGTCTGTGGAAGTC-3′; J2KI-2: 5′-CGTGGAGAGTCTGTAAGGCTCAA-3′), the floxed allele (Neo-Exc-LT: 5′-GCAAAGGGAGACAAGAAACGT-3′; Neo-Exc-RT: 5′-GACCAGTTGCTCCAGGGTTA-3′), the excised allele (Neo-Exc-RT; J2KI-4: 5′-TCACAAGCATTTGGTTTTGAAT-3′), and Cre recombinase (CRE1: 5′-CGCAAGAACCTGATGGACAT-3′; CRE2: 5′-TGCTGTCACTTGGTCGTGG-3′). Reactions were performed in 25 μL volumes containing 10× PCR buffer, 0.5 μL 10 mM dNTP mix, 1 μL primer pair (10 mM each), 0.2 μL polymerase, nuclease-free water, and 2 μL genomic DNA template. Thermal cycling conditions for the Geno, Flox, and Exc assays consisted of an initial denaturation at 95°C for 5 min, followed by 34 cycles of 95°C for 30 s, 57.5°C for 30 s, and 72°C for 40 s, with a final extension at 72°C for 5 min; Cre amplification was performed on diluted Geno/Flox/Exc PCR product using 21 cycles with an annealing temperature of 51.7°C. Amplicons were resolved on 2% agarose gels containing ethidium bromide alongside a low-range DNA ladder (SM1193; ThermoFisher Scientific) and visualized under UV illumination to determine genotypes based on expected band sizes for wild-type, heterozygous, and homozygous knock-in alleles. R26-M2rtTA;Col1a1-tetO-H2B-GFP compound mutant mice (JAX#016836) were used to allow doxycycline-inducible, fluorescent labelling of cells (Foudi et al., 2009). All mouse experiments were conducted in accordance with the National Institutes of Health Guide for the Care and Use of Laboratory Animals and were approved by the Institutional Animal Care and Use Committee at the University of California.

### Flow cytometry antibody staining of bone marrow (BM) hematopoietic cells

Total BM cells were isolated from mice by flushing the femur and tibia with staining medium composed of DPBS containing 2% heat-inactivated fetal bovine serum (FBS; FB-02; Omega Scientific) through a 0.7μm cell strainer (22363547; Fisherbrand). BM cells were pelleted, and then red blood cells were lysed using 1x ACK lysis buffer. For HSPC specific experiments, BM cells were lineage-depleted using MagniSort Mouse Hematopoietic Lineage Depletion kit (8804682974; Invitrogen) as per manufacturer’s instructions. Cells were incubated with the following anti-mouse antibodies: Pacific Blue lineage cocktail (containing: CD3, Ly-6C, CD11b, CD45R, Ter-119) (133310; Biolegend), APC/Fire-750 cKit (105838; Biolegend), BV605 Sca-1 (108134; Biolegend). For LT-HSC specific experiments cells were additionally incubated with APC CD135 (135310; Biolegend), PE CD150 (115904; Biolegend) and PE/Cyanine7 CD48 (103422; Biolegend). For myeloid-restricted progenitor subpopulation experiments, cells incubated with HSPC cocktails were additionally incubated with APC CD34 (17034182; eBiosciences), PE CD16/CD32 (553145; BD Pharmingen) and PE/Cyanine7 CD127 (25127182; eBiosciences). All antibodies were used at a 1:500 dilution factor for staining protocols. All cell isolations were performed using BD FACSAria Fusion sorter at the UCI Institute of Immunology Flow Cytometry Facility. All analyses of BM cells were performed on NovoCyte 3000. Data collection was performed using NovoExpress (v1.6.2) and analysis was performed using FlowJo (v10.10).

### In-vivo bromodeoxyuridine (BrdU) incorporation assay

Wild-type C57BL/6J mice were intraperitoneally injected with 5μg LPS (L2880; Sigma-Aldrich) with or without 100μg IL-10R blocking antibody (112702; BioLegend), or 1x PBS (DPBS; 14200075; ThermoFisher Scientific) for 72 or 24 hours prior to BM harvest for acute exposure analysis. All mice were injected with BrdU 16 hours prior to bone marrow harvest. For *in vivo* BrdU incorporation the above LT-HSC staining protocol was used but omitted APC-CD135 as APC BrdU Flow kit (552598; BD Pharmingen) was used to identify cycling cells (BrdU^+^) cells according to manufacturer’s protocol.

### H2B-GFP cell cycling assay

Doxycycline inducible R26-M2rtTA;Col1a1-tetO-H2B-GFP compound mutant mice were treated with 6.6g/L doxycycline (446061000; ThermoScientific) and 2g/L sucrose (A15583.0E; ThermoScientific) containing water for 4 weeks to ensure maximum GFP expression. After doxycycline treatment, mice were intraperitoneally injected with 5μg LPS (L2880; Sigma-Aldrich) with or without 100μg IL-10R blocking antibody (112702; BioLegend), or 1x PBS (DPBS; 14200075; ThermoFisher Scientific) for 2 or 4 weeks, 3 times weekly, prior to BM harvest for chronic exposure analysis. BM cells were used for long-term cell cycle analysis of LT-HSC populations after chronic and were subsequently stained using the LT-HSC protocol aforementioned, with the FITC channel being reserved for H2B-GFP assessment.

### Competitive transplantation assay

C57BL/6J mice were used as recipients for transplantation, mice were sex and age matched for transplants. Whole bone marrow of 1:1 equivalent LT-HSC from each competing donor was quantified using the above LT-HSC subpopulation antibodies and analyzed on the NovoCyte 3000 flow cytometer (Agilent). Recipients were lethally irradiated using an X-ray irradiator at 850cGy and rested for 24 hours before transplantation. Retroorbital transplantation was preformed using 1:1 equivalent LT-HSCs for a total of approximately 2 million BM cells from donor competitors into irradiated recipients. Chimerism for donor derived population longitudinal tracking was performed by saphenous vein bleeding collecting 20uL of blood followed by lysis of red blood cells using 1x ACK lysis buffer. Samples were then stained with FITC anti-CD45.1 (110706; Biolegend) and APC CD45.2 antibody (109814; Biolegend).

### Image Flowcytometry polarity assay

For image flowcytometry (IFC), BM cell isolation was performed and stained using the HSPC staining protocol aforementioned, followed by intracellular staining of alpha-tubulin using BD Cytofix/Cytoperm (512090KE; BD Biosciences) reagents as per manufacture’s recommendations (BD Biosciences). Alpha-tubulin was stained with a FITC conjugated antibody (627906; Biolegend). IFC was performed on an Amnis ImageStream MkII (Cytek Biosciences) at the UCI Flow Cytometry Facility, at 60X magnification at slow speed for highest sensitivity. For each sample, 50,000 single and focused cells were imaged for data acquisition. Data was analyzed using IDEAS® image analysis software (v6.2; Cytek Amnis) and 200 HSPC from each sample was manually counted and assessed for alpha-tubulin polarity.

### Quantification of peripheral blood cells, platelet counts, white blood cells count, histology and HSCs in the bone marrow

Mouse FITC anti-CD45.1 (110706; Biolegend) and APC CD45.2 antibody (109814; Biolegend). were used to quantify hematopoietic cell population. Scil Vet ABC hematology analyzer (Scil Animal Care Company) was used to obtain the complete blood cell count and platelet count from mouse peripheral blood. Upon termination of the transplant, spleens and femurs of transplanted mice were collected and fixed in formalin. Bone marrow and spleen samples were delivered to the UCI Experimental Tissue Resource (ERT) core for hematoxylin and eosin (H&E) staining.

### Bulk RNA-Sequencing

Wild-type C57BL/6J mice were intraperitoneally injected with 5μg LPS (L2880; Sigma-Aldrich) with or without 100μg IL-10R blocking antibody (112702; BioLegend), or phosphate buffered saline (DPBS; 14200075; ThermoFisher Scientific) for 72 or 24 hours prior to BM harvest for cell sorting using BD FACSAria Fusion sorter at the UCI Institute of Immunology Flow Cytometry Facility. Cells were sorted, using the HSPC staining protocol aforementioned, into RNA lysis buffer (R1060; Zymo-Research) and immediately processed using Zymo-Research Quick-RNA MiniPrep kit for RNA isolation as per manufacture’s protocol (R1058; Zymo-Research). Samples were stored at -20°C until sequencing at the UCI Genomics Research and Technology HUB. Fastq data files were converted to BAM files using HISAT2 v2.2.0 and aligned against GRCm38 genome using default settings (Kim et al., 2019). Raw counts were extracted from BAM files using Subread function featureCounts v2.1.1 under default settings, probe counts were saved onto a csv file. An associated meta data csv file was created for differential analysis. Differentially expressed genes (DEGs) were elucidated in R v4.3.2 using DESeq2. Single sample Gene Set Enrichment Analysis was done in R using the Gene Set Variation Analysis (GSVA) package v3.21. Plots were generated in RStudio. Gene set enrichment analysis (GSEA) was performed the GSEA v4.3.3 software. Refer to *Data Availability* section for more information on the bioinformatics used.

### Statistical analysis

Data are represented as means ± standard deviations (S.D.), or as means ± standard error of the mean (SEM) for blood chimerism following transplantation assays. GraphPad Prism was used for statistical analysis for all other data. Dots on box and bar plots represent biological replicates. In line plots, biological replicates are noted on the figure or legend. Two-tailed unpaired *t-*test was used when two groups were compared, with Holm-*Śídák* correction when multiple comparisons were performed. One-way ANOVA was used when three or more groups were compared, with Tukey’s correction when multiple comparisons were performed. Two-way ANOVA was used when three or more groups with two independent variables were compared, with Tukey’s or *Śídák* correction when multiple comparisons were performed. R studio, and Gene-Set Enrichment Analysis was used for RNA-sequencing data analysis and associated plot curation. Statistical significance was determined by Wald test with Benjamini–Hochberg FDR correction as standard using DESeq2. Statistical significance is indicated in figures or figure legends either by exact p-values or by symbols as follows: p < 0.05 (*), < 0.005 (**), < 0.0005 (***), and < 0.0001 (****). Absence of p-value or symbol denotes statistical non-significance unless otherwise specified. Data collection and analysis were not performed blind to the conditions of the experiments.

### Online supplemental material

Figure S1 provides information to the flowcytometry gating schemes for HSPC, LT-HSC, and the myeloid and lymphoid progenitors, including BrdU and H2B-GFP compositions. Figure S2 provides additional information accompanying the bulk cell RNA-seq results in Figure 2. Figure S3 provides longitudinal chimerism gating schemes to Figures 4 and 6, additionally including accompanying pre-exposure transplants to Figure 4. Figure S4 provides additional information for BrdU assays of HSPCs alongside Figure 5 and further contains additional information for WT:WT transplant, and the repeated VF:WT transplant to accompany Figure 6.

### Data availability

The RNA-seq data that support the findings of this study have been deposited in Gene Expression Omnibus under the accession code <u>GSE308698</u>. The genome used for alignment was *Mus musculus* genome GRCm38 and is available at https://www.ncbi.nlm.nih.gov/datasets/genome/GCF_000001635.20/. R-script used for DEG analysis and gene ontology can be obtained at https://github.com/wwadley-lucas/bulkRNA-seq-script-R. Any other data are available from the corresponding author upon reasonable request.

## Acknowledgments

We foremost acknowledge the University of California Irvine’s shared resources whereby research reported in this publication was supported by the National Cancer Institute of the National Institutes of Health under award number P30CA062203. This work utilized resources of the UCI Genomics Research and Technology Hub (GRT Hub) parts of which are supported by NIH grants to the Comprehensive Cancer Center (P30CA-062203) and the UCI Skin Biology Resource Based Center (P30AR075047) at the University of California, Irvine, as well as to the GRT Hub for instrumentation (1S10OD010794-01and 1S10OD021718-01. The authors also wish to acknowledge the support of the Chao Family Comprehensive Cancer Center IFI Flow Cytometry Facility Shared Resource, and the Chao Family Comprehensive Cancer Center Experimental Tissue Shared Resource supported by the National Cancer Institute of the National Institutes of Health under award number P30CA062203. The content is solely the responsibility of the authors and does not necessarily represent the official views of the National Institutes of Health.

This research was funded through grants from the National Institute of Diabetes and Digestive and Kidney Diseases (NIDDK) and National Cancer Institute (NCI), including R01DK136069 and R37CA271172, respectively, to Angela Fleischman. This study was also funded in part by the National Heart Lung and Blood Institute predoctoral grant F31HL176174 to William Lucas Wadley. This study was also funded in part by the University of California Irvine Institute of Immunology training grant T32AI0602573 to H.Y. Lai.

## Author contributions

William Lucas Wadley: Conceptualization, Data curation, Formal analysis, Investigation, Methodology, Resources, Validation, Writing - original draft, Writing – review and editing. H. Huang: Investigation, Methodology, Validation, Writing - review and editing. H. Y. Lai: Conceptualization, Data curation, Formal analysis, Investigation, Validation, Writing - review and editing. J. Chen: Investigation, Writing – review and editing. J. Heidmann: Investigation. E. Soyfer: Investigation, Methodology. K. Guillermo: Investigation, Resources. E. Arora: Investigation. L. Chen: Investigation. B. Hoover: Investigation, Formal analysis, Methodology. A. Fleischman: Conceptualization, Funding Acquisition, Methodology, Project Administration, Resources, Supervision, Validation, Writing - original draft, Writing – review and editing.

---

**Supplemental Figure 1.**
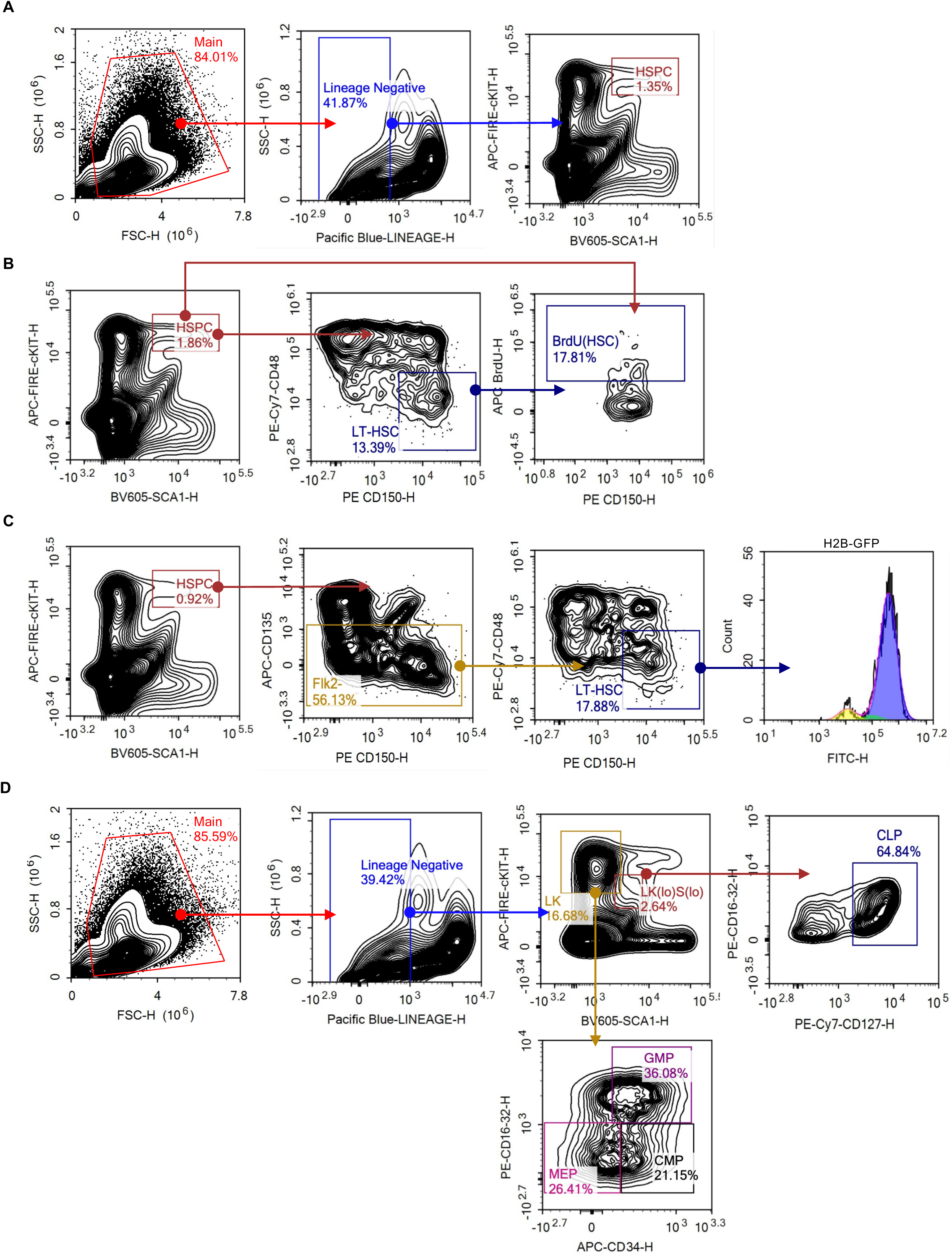
Flow cytometry gating for. (A) HSPC’s defined as Lineage negative, cKIT+, Sca-1+. (B) LT-HSC’s regarding BrdU incorporation defined as CD150+, CD48-of the HSPC group, commonly referred to as LKS-SLAM. BrdU gated on the APC channel and as per manufacturer’s instructions. (C) LT-HSCs used normally or in H2B-GFP analysis defined as Flk2- (CD135-), CD150+, CD48- of the HSPC group, with H2B-GFP being assessed in the FITC channel. (D) Myeloid restricted progenitors defined as Lineage negative, cKIT+, Sca-1- (LK) and the subpopulations common myeloid progenitors (CMPs; CD34^high^/CD16/32^low^), granulocyte–macrophage progenitors (GMPs; CD34^+^/CD16/32^+^), and megakaryocyte–erythroid progenitors (MEPs; CD34^−^/CD16/32^low^). Common lymphoid progenitors were defined as CD127+ of the Lineage negative, c-KIT^low^, and Sca-1^low^.

**Supplemental Figure 2.**
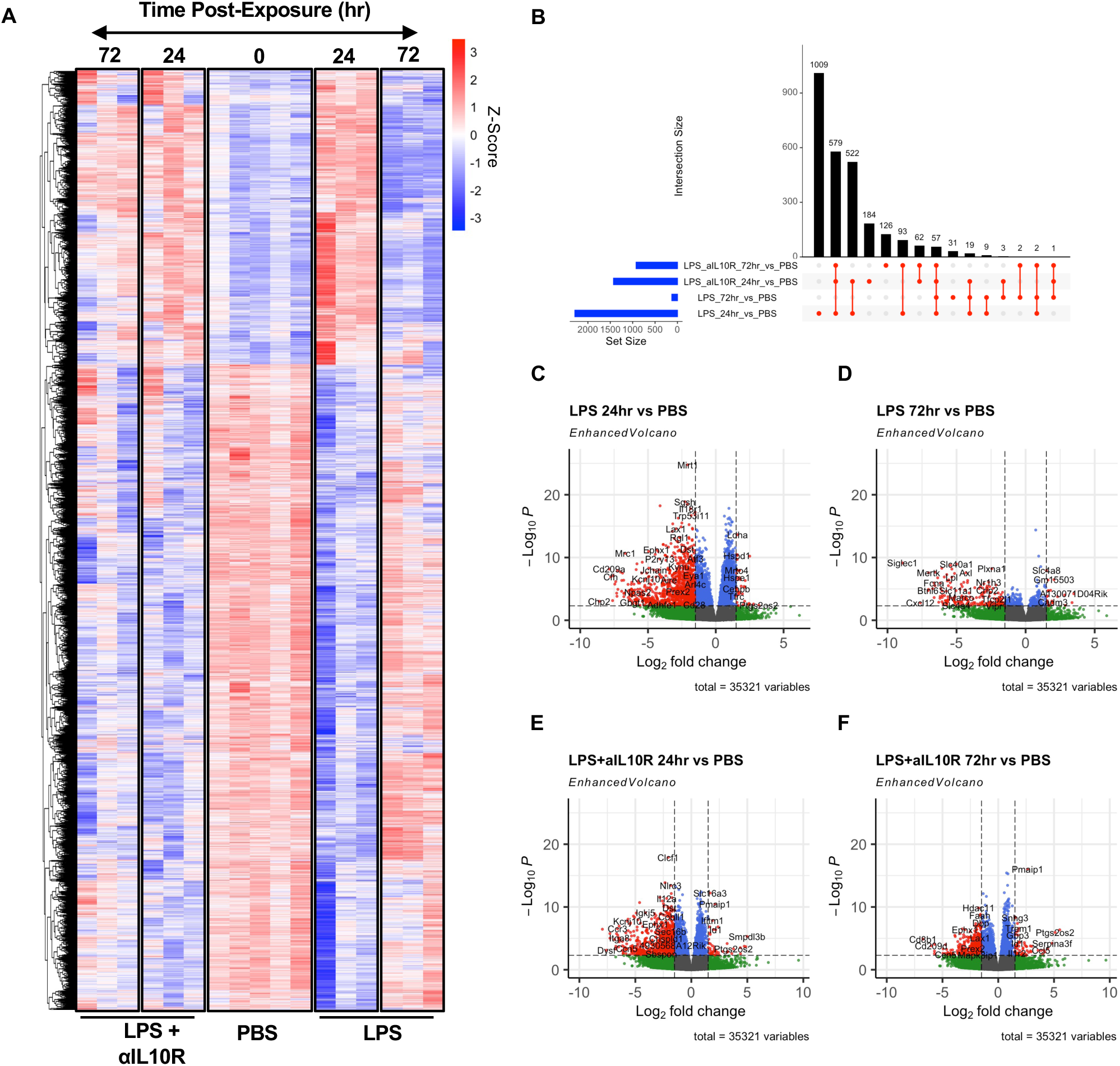
Bulk RNA Seq supporting data. (A) Heatmap of all differential expressed genes (DEGs) assessed via DESeq2. (B) Upset plot of bulk RNA seq depicting shared DEGs. (C-F) Volcano plots of bulk RNA seq data all compared to PBS Day 0 control group. Statistical significance was determined by Wald test with Benjamini–Hochberg FDR correction as standard using DESeq2.

**Supplemental Figure 3.**
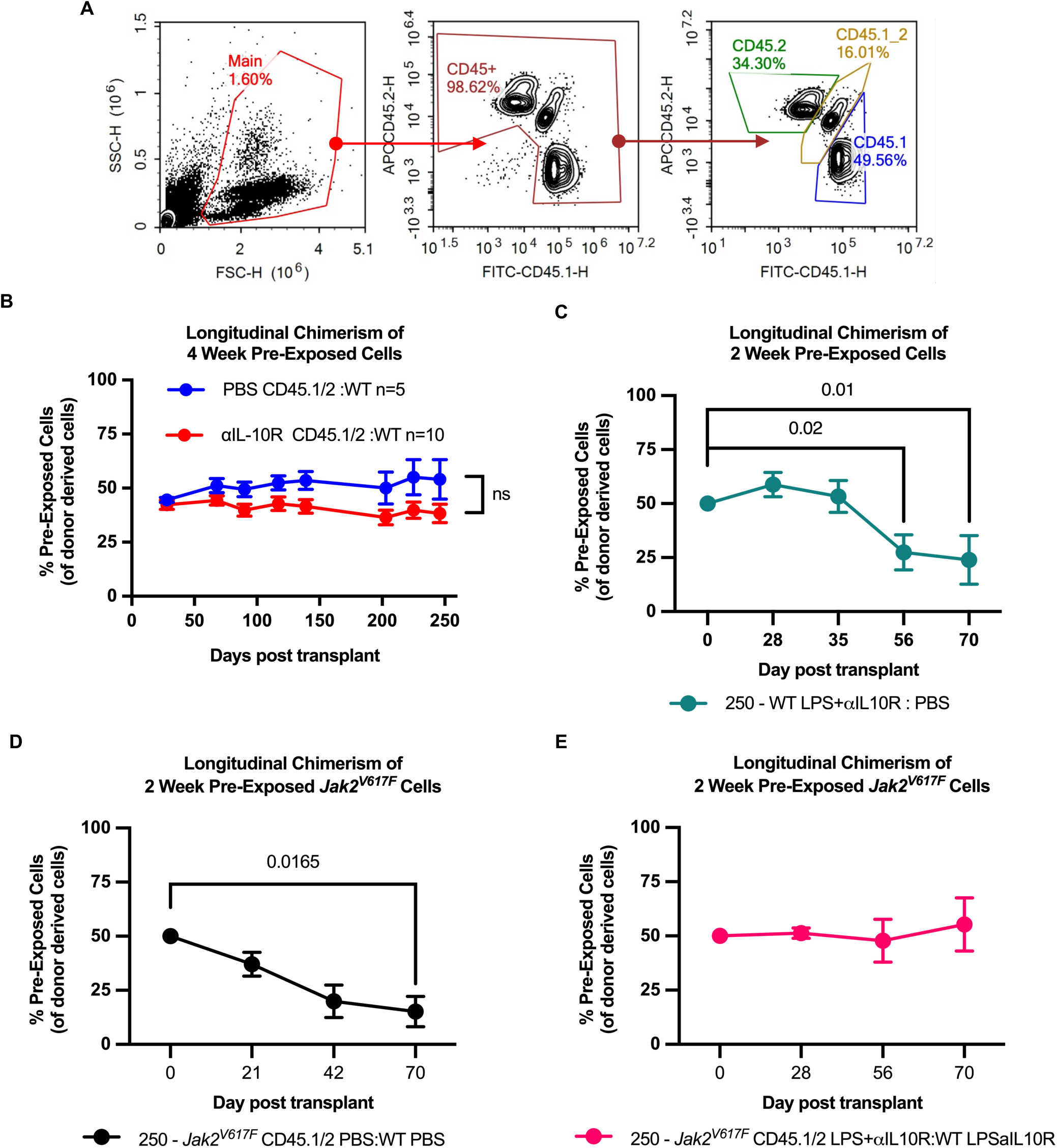
(A) Flowcytometry gating for longitudinal chimerism assessment of transplants (CD45.1, CD45.2, CD45.1/2) from peripheral blood draws. (B) Pre-exposure competitive transplant assay of IL-10R blocking antibody alone, compared against PBS control pre-exposure, all data points were found to be non-significant. (C-E) HSC sorted competitive transplant from pre-exposed mice from LPS and anti-IL-10R blockade. Supportive data for Figures 4 and 6’s transplant data with whole bone marrow to show outgrowth or decline in competition correlates to HSC specific competition transplantation data, n = 5 for all conditions, the “250” notation on each plot indicates 250 HSCs of each competitive group were used for transplantation. Statistical significance was determined by repeated measures (RM) two-way ANOVA (C-G). If not specifically indicated, P-value < 0.05, 0.005, 0.0005, and <0.0001 are represented as *, **, ***, and **** respectively. Notation of ns indicates no statistical significance.

**Supplemental Figure 4.**
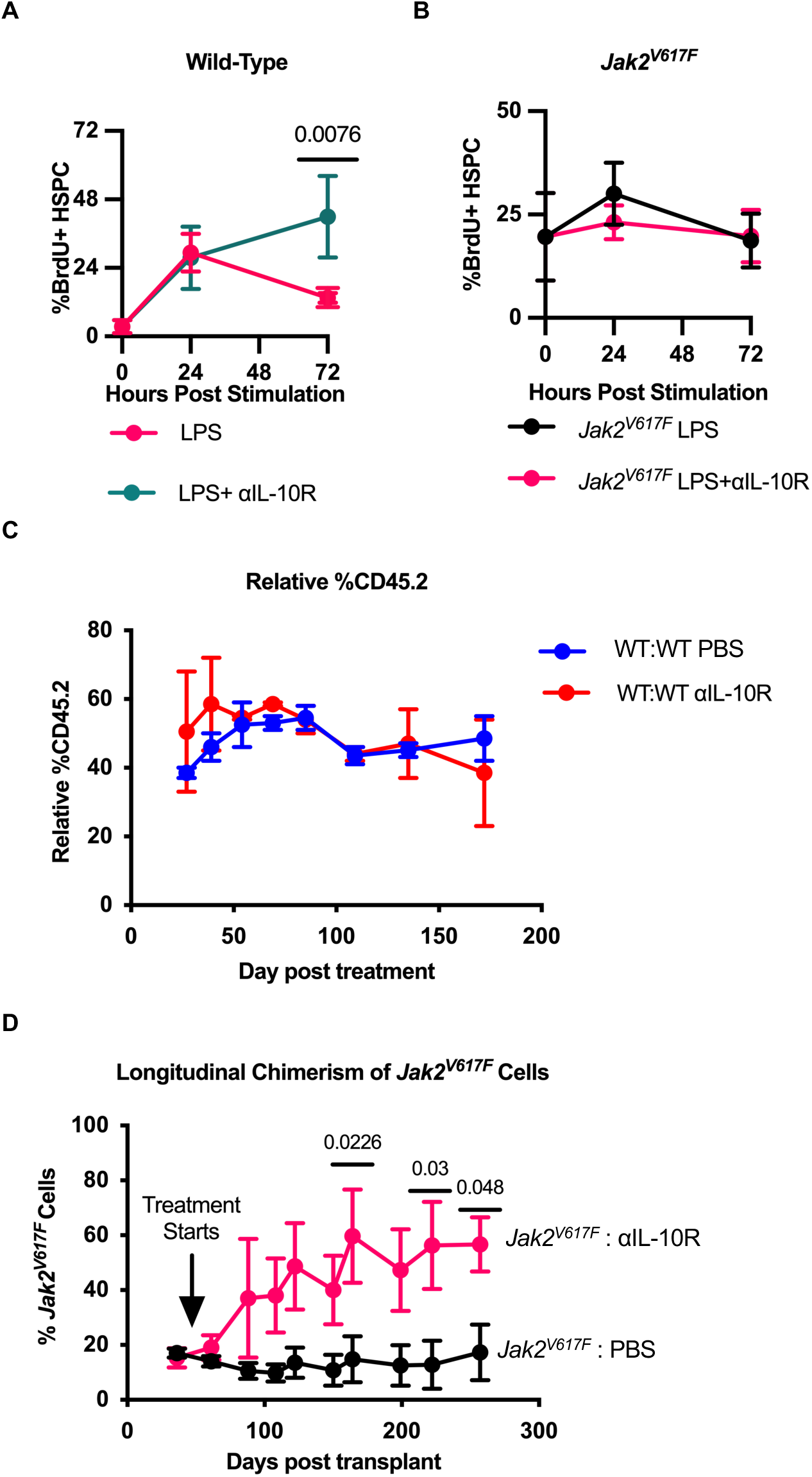
(A) BrdU quantification of WT HSPCs from acute exposure with LPS and IL-10R blocking antibody (n = 3 for all conditions and groups), SD error bars presented. (B) BrdU quantification of *Jak2^V617F^*HSPCs (*right*) from acute exposure with LPS and IL-10R blocking antibody (n = 3 for all conditions and groups), SD error bars presented. (C) CD45.2 chimerism of WT:WT control competitive transplantation and starting a blocking IL-10R antibody treatment (n = 2) 60-days post-transplantation, or no treatment control (n = 2). SEM error bars presented. (D) *Jak2^V617F^* chimerism and starting a blocking IL-10R antibody treatment (n = 3) 49-days post-transplantation, or no treatment control (n = 4). SEM error bars presented. Statistical significance was determined by repeated measures (RM) two-way ANOVA with *Śídák* correction(C and D). If not specifically indicated, P-value < 0.05, 0.005, 0.0005, and <0.0001 are represented as *, **, ***, and **** respectively.

